# Novel roles for the LRRC56 protein, an IFT cargo protein, in docking of dynein arms in *Trypanosoma brucei*

**DOI:** 10.1101/2023.08.09.552631

**Authors:** Serge Bonnefoy, Aline Araujo Alves, Eloïse Bertiaux, Philippe Bastin

## Abstract

Outer dynein arms (ODAs) are responsible for ciliary beating in eukaryotes. They are assembled in the cytoplasm and shipped by intraflagellar transport (IFT) before attachment to microtubule doublets via the docking complex. The LRRC56 protein has been proposed to contribute to ODAs maturation. Mutations or deletion of the *LRRC56* gene lead to reduced ciliary motility in all species investigated so far, but with variable impact on dynein arm presence. Here, we investigated the role of LRRC56 in the protist *Trypanosoma brucei,* where its absence results in distal loss of ODAs, mostly in growing flagella. We show that LRRC56 is a transient cargo of IFT trains during flagellum construction and surprisingly, is required for efficient attachment of a subset of docking complex proteins present in the distal portion of the organelle. This relation is interdependent since the knockdown of the distal docking complex prevents LRRC56’s association with the flagellum. Intriguingly, *lrrc56^-/-^* cells display shorter flagella whose maturation is delayed. Inhibition of cell division compensates for the distal ODAs absence thanks to the redistribution of the proximal docking complex, restoring ODAs attachment but not the flagellum length phenotype. This work reveals an unexpected connection between LRRC56 and the docking complex.

## Introduction

Ciliary motility is driven by the action of outer (ODA) and inner (IDA) dynein arms, two large multi-protein complexes attached to the A-tubule of axonemal doublets. ODA and IDA are usually present on all 9 doublet microtubules and distributed all along the axoneme. They are composed of a variable number of heavy, intermediate and light chains (for a review on dynein composition, see King, 2016). Only the heavy chains have motor activity while intermediate and light chains are essential for the assembly and regulation of the complex. Dynein arms are assembled in the cytoplasm by a specific machinery itself composed of several subunits (Omran *et al*., 2008). They are then transported by intraflagellar transport (IFT) (Dai *et al*., 2018) and finally attached to the microtubule via the docking complex, a little structure itself fixed to the A-tubule (Qiu and Roy, 2022).

This machinery is remarkably well conserved throughout evolution and defects in any component of this sophisticated pathway perturbs dynein arm assembly, resulting in altered motility in all organisms investigated so far. This includes humans, where mutations in genes involved in dynein arm composition, assembly or transport cause Primary Ciliary Dyskinesia (PCD, MIM: PS244400). This disease is characterised by impaired mucociliary clearance and increased susceptibility to respiratory infections, infertility and *situs inversus* or other laterality disorders (Legendre *et al*., 2021).

Model organisms such as the green alga *Chlamydomonas* have been instrumental in deciphering the composition, assembly and function of dynein arms (Silflow and Lefebvre, 2001). Identification of the first *PCD* gene that encodes the intermediate dynein chain 1 (DNA1 or IC78), a central component of the outer dynein arm, was guided by exhaustive structural and genetic knowledge in the green alga (Pennarun *et al*., 1999). Later on, it turned out that many components of the dynein arms and their assembly machinery were largely conserved in eukaryotes with motile cilia.

However, an unusual situation has been recently reported for the *LRRC56/ODA8* gene, which encodes a protein containing multiple leucine-rich repeats (Leucine-Rich Repeat Containing 56 protein) that is conserved in organisms with motile cilia assembled by IFT (Desai *et al*., 2015). Its impairment either by mutations in human patients or *Chlamydomonas*, or by double gene deletion in *T. brucei* revealed a clear contribution to ciliary beating in all three species but surprisingly, the ultrastructural consequences were distinct (Bonnefoy *et al*., 2018). The *Chlamydomonas* null mutant is termed *oda8* and is characterised by the absence of outer dynein arms all along the axoneme (Kamiya, 1988; Desai *et al*., 2015). By contrast, a predicted null mutation in a human patient is not accompanied by visible structural defects in dynein arms of cilia from nasal biopsy samples, despite a dyskinetic beat pattern observed in air-liquid interface culture (Bonnefoy *et al*., 2018). Even more intriguingly, the knockout of *LRRC56* in *T. brucei* results in the formation of flagella where ODA is lacking from only the distal half of the axoneme, mostly in growing flagella (Bonnefoy *et al*., 2018).

In *Chlamydomonas*, the precise localisation of the LRRC56 protein could not be determined by immunofluorescence but cell fractionation revealed that an HA-tagged version of the protein was present mostly in the cytoplasm and in the matrix of the flagellum (Desai *et al*., 2015), a distribution similar to that of IFT proteins (Ahmed *et al*., 2008). Using *in vivo* imaging, IFT-mediated transport of outer dynein arm components has been demonstrated upon expression of the intermediate chain IC2 (also called DNAI2) fused with a mNeonGreen (mNG) fluorescent reporter. In the *oda8/lrrc56* mutant, the frequency of ODA transport events is dramatically reduced, indicating defects either for entry in the flagellum or for association with IFT trains (Dai *et al*., 2018). Pull-down experiments in HEK293 cells showed that the human LRRC56 co-immunoprecipitates with IFT88, also suggesting an association with IFT (Bonnefoy *et al*., 2018). In trypanosomes, a YFP fusion protein expressed *in situ* from the *LRRC56* locus was found in the matrix of the flagellum, mainly at the distal portion of growing flagella and in an IFT-dependent manner. All these data indicate that LRRC56 interacts somehow with the IFT machinery in a wide range of organisms, but its precise functions remain to be deciphered, as well as why the absence of LRRC56 leads to unusually variable structural phenotypes between different organisms.

In an effort to explain the trypanosome situation where the *lrrc56* knockout mutant only lacks dynein arms in the distal portion of its axoneme (and yet mostly during elongation and not so much in mature flagella), we considered two main hypotheses. First, the axoneme is heterogeneous all along its length (Subota *et al*., 2014; Edwards *et al*., 2018; Billington *et al*., 2022; Billington *et al*., 2023). In particular, there are two distinct docking complexes: a proximal one that covers about the first half of the flagellum and a distal one, present only on the other half. Each is made of at least two subunits that contain coiled-coil domains (proximal: pDC1 and pDC2, distal: dDC1 and dDC2). pDC1 and dDC1 are related to the CCDC151/ODA3 family (Koutoulis *et al*., 1997; Hjeij *et al*., 2014; Jerber *et al*., 2014) whilst pDC2 and dDC2 are related to CCDC114/ODA1 (Takada *et al*., 2002; Knowles *et al*., 2013; Onoufriadis *et al*., 2013). In this context, LRRC56 could only be required for the addition of dynein arms to the distal docking complex, but this hypothesis would not explain why this is mostly encountered in growing flagella. Second, dynein arms are added to growing microtubules. If LRRC56 was required for efficient entry, transport or maturation of dynein arms in the trypanosome flagellum as observed in *Chlamydomonas* (Dai *et al*., 2018), one could imagine that the rate of addition of dynein arms in *lrrc56^-/-^*cells is too slow relative to that of elongation of the axoneme, hence limiting ODA presence to the proximal portion of the flagellum. This is supported by the fact that in the *lrrc56* knockout mutant, the mature flagella possess a higher proportion of dynein arms than the growing flagella (Bonnefoy *et al*., 2018).

We show here that LRRC56 is an IFT cargo and that a combination of the two proposed models can explain the distal absence of dynein arms in growing flagella and its partial compensation in mature ones but in a rather unexpected manner. In the absence of LRRC56, proteins of the distal docking complex are present in very low concentration in the middle portion of the flagellum and absent from its distal part, hence explaining the lack of dynein arms in that region in growing flagella, and shedding light on an unexpected connection between LRRC56 and the docking complex. This appears interdependent as the knockdown of dDC1 or dDC2 prevents LRRC56 from reaching the flagellum. As flagella age over time, the proximal docking complex occupies more extensive territories than in control cells, allowing further association of dynein arms that almost reach the distal end. These results answer the question as to why dynein arms were missing only in the distal portion of young flagella and how it is compensated as flagella mature.

## Material and methods

### Trypanosome cell culture

Cells used for this work were derivatives of *T. brucei* strain Lister 427 and were cultured in SDM79 medium supplemented with hemin and 10% fetal calf serum (Brun and Schönenberger, 1979).

### Expression of endogenous tagged *T. brucei* proteins

N-terminus tagging of LRRC56 with mNeonGreen was carried out as described (Dean *et al*., 2015), using p2675 derived template plasmid (Kelly *et al*., 2007) to amplify the puromycin drug resistance cassette as well as the gene encoding the mNeonGreen fluorescent reporter using forward primers containing 80 nucleotides of the 5′ UTR of *LRRC56* (Tb927.10.15260) followed by the 20 nucleotides plasmid primer binding sequence and reverse primers consisting of the first 80 nucleotides of the target *lrrc56* open reading frame (Dean *et al*., 2015) in reverse complement orientation, followed by the last 20 reverse complement nucleotides of the mNeonGreen gene without the stop codon. Endogenous N-terminus tagging of *IFT81* (a member of the IFT-B complex) *(Bhogaraju et al., 2013; Kubo et al., 2016)* with tandem Tomato (tdT) fluorescent reporter (Shaner *et al*., 2004) was carried out using p2845 derived plasmid to amplify the blasticidin drug resistance cassette as well as the Tandem Tomato gene using forward primers containing 80 nucleotides of the 5′ UTR of IFT81 (Tb927.10.2640) followed by the 20 nucleotides plasmid primer binding sequence and reverse primers consisting of the first 80 nucleotides of the target *IFT81* open reading frame (Dean *et al*., 2015) in reverse complement orientation, followed by the last 20 reverse complement nucleotides of the *tdT* gene without the stop codon.

For the generation of the mNG::dDC1 and mNG::dDC2 expressing cell lines, the first 750 nucleotides of dDC1 (Tb927.5.1900) and dDC2 (Gene DB number Tb927.11.16090) without the ATG were chemically synthesized (GeneCust) and cloned in frame with the mNG gene (Shaner *et al*., 2013) within the HindIII and ApaI sites of p2675 vector (Kelly *et al*., 2007) to generate p2675mNGdDC1 and p2675mNGdDC2. Before nucleofection, plasmid constructs were linearized within the dDC1 and dDC2 sequence with the enzyme ClaI and XcmI, respectively. For the generation of the mNG::pDC1 cell lines, the p2675mNGdDC2 plasmid was used as a template to amplify the puromycin drug resistance cassette as well as the gene encoding the mNeonGreen fluorescent reporter and using forward primers containing 80 nucleotides of the 5′ UTR of pDC1 (Tb927.8.4400) followed by the 20 nucleotide plasmid primer binding sequence and reverse primers consisting of the first 80 nucleotides of the target pDC1 open reading frame (Dean *et al*., 2015) in reverse complement orientation, followed by the last 20 reverse complement nucleotides of the mNeonGreen gene without the stop codon. For the generation of the tdT::pDC1 cell line, p2845tdTIFT140SAT plasmid was obtained from p2845tdTIFT140 plasmid (Bertiaux *et al*., 2018b) by replacing the selectable marker for blasticidin resistance with the SAT-1 streptothricin acetyltransferase gene (Joshi *et al*., 1995) which confers resistance to nourseothricin. p2845tdTIFT140SAT was used as a template to amplify the SAT resistance cassette as well as the Tandem Tomato gene (Shaner *et al*., 2004) using forward primers containing 80 nucleotides of the 5′ UTR of pDC1 followed by the 20 nucleotide plasmid primer binding sequence and reverse primers consisting of the first 80 nucleotides of the target pDC1 open reading frame (Erdmann *et al*., 2019) in reverse complement orientation, followed by the last 20 reverse complement nucleotides of the *tdT* gene without the stop codon.

Sterile DNA was obtained following ethanol precipitation, and quantification was performed with a nanodrop. Transfection in the appropriate cell lines was achieved by nucleofection using program X-014 of the AMAXA Nucleofector® apparatus (Lonza) as previously described (Burkard *et al*., 2007) with 10 *µ*g linearized plasmids or 5 *µ*g PCR product for homologous recombination with the target allele (Dean *et al*., 2015). Transgenic cell lines were obtained following appropriate drug selection according to the recipient cells used, then cloned if required by serial dilution.

### Generation of cell lines for RNAi knockdown

The 2913 cell line expressing the T7 RNA polymerase and the tetracycline repressor has been described previously (Wirtz *et al*., 1999). For generation of the *dDC2^RNAi^* and *dDC1^RNAi^* cell lines, a 498 bp fragment of *dDC2* (Tb927.11.16090) and a 505 bp fragment of *dDC1 (*Tb927.5.1900), both flanked by HindIII (upstream) and XhoI (downstream) sites, were selected using the RNAit algorithm (http://dag.compbio.dundee.ac.uk/RNAit/), to ensure that the fragment lacked significant identity to other genes and to avoid cross-RNAi (Redmond *et al*., 2003). These fragments were generated by chemical synthesis by GeneCust Europe (Boynes, France) and cloned into the corresponding HindIII-XhoI site of the digested pZJM vector (Wang *et al*., 2000) allowing for tetracycline-inducible expression of dsRNA generating RNAi upon transfection in the 2913 recipient cell line. The dsRNA is expressed from two tetracycline-inducible T7 promoters facing each other in the pZJM vector.

### Intraflagellar trafficking analysis

Double-tagged mNG::LRRC56 and tdT::IFT81 *lrrc56*^-/-^ cells were used to image two color fluorescence simultaneously in live cells. Live cells were spread on a glass slide, covered with a coverslip and immediately observed with a spinning-disk confocal microscope (UltraVIEW VoX; Perkin Elmer) equipped with an oil-immersion objective Plan Apochromat 100×/1.57 NA (ZEISS). Videos were acquired using Volocity software with two EM-CCD cameras (ImagEM X2 C9100-23B, Hamamatsu) in streaming mode. The samples were kept at 27°C using a temperature-controlled chamber. Sequences of 15 s were acquired with an exposure time of 100 ms per frame in dual-color imaging mode. Kymographs were extracted and analyzed with Icy software (de Chaumont et al., 2012) using the plugin Kymograph Tracker v2.1.3, as described (Chenouard *et al*., 2010; de Chaumont *et al*., 2012; Buisson *et al*., 2013). For kymograph extraction, the tdT::IFT81 signal was used as a reference to determine the region of interest. That same region of interest was applied to the mNG::LRCC56 signal of the corresponding cell.

After kymograph extraction, the Icy software was used for the image background subtraction of green and red anterograde kymographs, followed by the generation of the Manders’ coefficients between the two signals with the Colocalization Studio plugin, using the same regions of interest as for the speed and frequency measurements.

### Indirect immunofluorescence assay (IFA)

Cultured trypanosomes were washed twice in SDM79 medium without serum and spread directly on poly-L-lysine coated slides (Thermoscientific, Menzel-Gläser), air dried, then fixed in methanol at −20°C for 5 min followed by a rehydration step in PBS for 15 min. For cytoskeleton analysis, cells were settled onto poly-L-lysine coated slides, and cytoskeletons were prepared as described (Bonhivers *et al*., 2008) by extraction with 0.4% NP-40 in 100 mM Pipes, pH 6.9, 2 mM EGTA, 1 mM MgSO_4_, and 0.1 mM EDTA. Following 2 washes in PIPES, cells were fixed in methanol at −20°C for 5 min and then rehydrated for 10 min in PBS. For immunodetection, slides were incubated for 1 h at 37°C with the appropriate dilution of the first antibody in 0.1% BSA in PBS: mAb25 against the axonemal protein TbSAXO1 (Dacheux *et al*., 2012) a mouse polyclonal antiserum against DNAI1 (Duquesnoy *et al*., 2009), or the 32F6 IgG2c monoclonal antibody against mNeonGreen protein (Chromotek). After several 5 min washes in PBS, species and subclass-specific secondary antibodies coupled to the appropriate fluorochrome (FITC, Alexa 488, Cy3 or Cy5, Jackson ImmunoResearch) were diluted 1/400 in PBS containing 0.1% BSA and were applied for 1 h at 37°C. After washing in PBS as indicated above, cells were stained with a 1 *µ*g/ml solution of the DNA-dye DAPI (Roche) and mounted with Slowfade antifade reagent (Invitrogen). Slides were either stored at −20°C or immediately observed with a DMI4000 microscope (Leica) with a 100X objective (NA 1.4) using Hamamatsu ORCA-03G or Prime95B Photometrics cameras with an EL6000 (Leica) as the light source. Image acquisition was performed using Micro-manager software and images were analyzed using ImageJ (National Institutes of Health, Bethesda, MD)

### Teniposide treatment

For inhibition of cell division, the topoisomerase II inhibitor teniposide (Sigma SML0609), was dissolved in DMSO and added to trypanosome cultures at a final concentration of 200 μM (Robinson and Gull, 1991; Bertiaux *et al*., 2018b) during 20 hours. As a control, an identical volume of DMSO was added to the flasks.

### Statistical Analysis

GraphPad Prism 8 application was used for all analyses. All data was analyzed using an unpaired t-test with Welch’s correction.

## RESULTS

### mNG::LRRC56 is transported by IFT

Multiple data indicate that LRRC56 is somehow related to IFT (Desai *et al*., 2015; Bonnefoy *et al*., 2018). To evaluate if LRRC56 is trafficking inside the flagellum and could be a cargo of IFT, we generated endogenous double-tagged mNG::LRRC56 and tdT::IFT81 (a classic IFT marker, (Fort *et al*., 2016; Bertiaux *et al*., 2018a) in a wild-type background to image two color fluorescence simultaneously in live cells. Acquisitions were performed on monoflagellated cells displaying an mNG::LRRC56 fluorescent signal, therefore corresponding to cells that inherited a new flagellum after cytokinesis, LRRC56 being absent from the mature flagellum. Video observation showed the expected trafficking of the mNG::IFT81 protein, with generally processive anterograde movement from base to tip (Figure 1Aa and Supplemental Movie S1) exhibiting the expected speed and frequency (Figure 1C, D). A few arrested trains were sometimes observed (Figure 1A and Supplemental Movie S1). These results agree with previously reported data for IFT protein trafficking in *T. brucei* (Huet *et al*., 2014; Fort *et al*., 2016; Bertiaux *et al*., 2018a). When observing the green channel, short fluorescent particles of mNG::LRRC56 displayed obvious anterograde movement (Figure 1Ac and Supplemental Movie S1). However, they appeared less processive than tdT::IFT81 and were mostly restricted to the distal portion of the flagellum (Figure 1Ac). Kymographs (space-time plots) analysis confirmed the presence of anterograde trafficking of tdT::IFT81 along the flagellum (Figure 1Ab) and revealed numerous traces of mNG::LRRC56 particles movement in the distal portion of the flagellum (Figure 1Ad). However, the pattern looked more complex than for mNG::IFT81, with a combination of arrested and moving particles, some showing alternation of movement with periods of stalling at various places along the distal portion of the flagellum (Figure 1Ad). The speed average of mNG::LRRC6 particles was similar to IFT trains but significantly slower (Figure 1C), while their frequency was higher (Figure 1D). Merging the kymographs for tdT::IFT81 and mNG::LRRC56 revealed a frequent association between the two proteins (Figure 1A, B), providing the first evidence that LRRC56 is a cargo of the IFT machinery (Figure 1Ae-f). When IFT trains were arrested, mNG::LRRC56 was also frequently associated with them (dotted white lines, Figure 1Af). However, the IFT and LRRC56 association appeared transitory, with several loading and premature unloading events (magenta and white lines, Figure 1Af). Intriguingly, many mNG::LRRC56 particles not associated with IFT trains were also observed (green lines, Figure 1Af) usually with lower speeds. Mander’s colocalization coefficient demonstrated that 36% of mNG::LRRC56 moving particles were colocalizing with tdT::IFT81 and that 40% of IFT trains were colocalizing with mNG::LRRC56 (Figure 1E). This shows that only a fraction of the mNG::LRRC56 flagellar pool is transported as cargo of the IFT system. Although most of the mNG::LRRC56 signal was restricted to the distal portion of the flagellum, some anterograde particles were detected in the proximal portion of the flagellum. A second cell is shown in Supplemental Figure S1 and Supplemental Movie S2.

**Figure 1.**
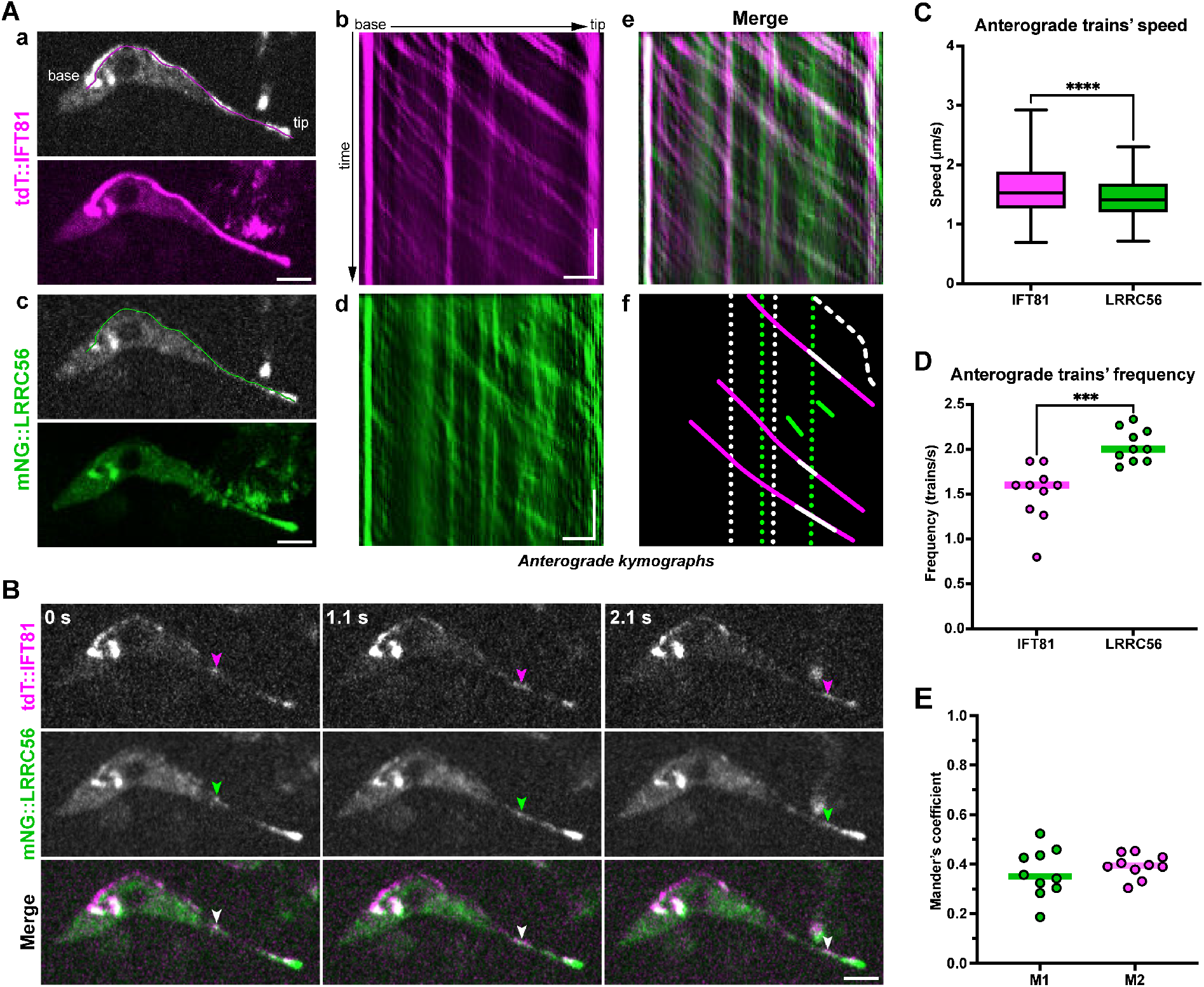
IFT transport of mNG::LRRC56 protein. (A) Kymograph analysis of a cell expressing tdT::IFT81 and mNG::LRRC56 following *in situ* tagging. (a) A still image of the red channel with tdT::IFT81 (top) shows this protein concentrates at the base of the flagellum and distributes along the length until the tip. In magenta, the region of interest used to extract the kymographs. Following the 15-second timelapse acquisition, the temporal projection of tdT::IFT81 (bottom) reveals its processive anterograde movement in the entire flagellar length. Scale bar: 3 *µ*m. (b) The IFT81 anterograde kymograph of the cell in (a) shows this protein displacing from base to tip with a few arrested trains. Horizontal scale bar: 3 *µ*m, vertical scale bar: 3 s. (c) A still image of the green channel (top) of the same cell in (a) showing mNG::LRRC56 (top). The same region of interest shown in green in (a) was used to extract the kymographs. The mNG::LRCC56 temporal projection (bottom) demonstrates that this protein concentrates at the distal portion of the flagellum. Scale bar: 3 *µ*m. (d) The anterograde kymograph of the cell in (c) shows mNG::LRRC56 either as arrested material or as particles moving from the base toward the tip. Horizontal scale bar: 3 *µ*m, vertical scale bar: 3 s. (e) Merge of IFT81 (b) and LRRC56 (d) anterograde kymographs reveals their partial colocalization. (f) A schematic summary of different events observed in (e). The dotted lines represent the arrested material of LRRC56 alone (green) or associated with IFT81 (white). The dashed line indicates a particle containing IFT81 and LRRC56 that alternates between moving and stalling. Continuous lines show IFT81 particles (magenta) carrying LRRC56 for a short part of their trajectory (white lines). LRRC56 is also moving without being associated with IFT (green lines). (B) Still images of the same cell in (A) at different time points confirm the displacement of an IFT particle (tdT::IFT81) transporting LRRC56 (arrowheads). See Supplemental Movie S1 for the full sequence. Images were processed to enhance contrast. Scale bar: 3 *µ*m. (C) Quantification of anterograde trafficking speed of mNG::IFT81 (magenta) (n=227 from 10 cells) and mNG::LRRC56 (green) (n=306 from 10 cells) in monoflagellated cells. **** p<0.0001. (D) Quantification of the frequency of anterograde particles (n=10 cells) shows LRRC56 particles are more abundant than IFT81. ***p=0.0004. (E) Manders’ colocalization analysis indicates a relatively low correlation between the trajectories of IFT81 and LRRC56.

### Partial absence of the distal docking complex in *lrrc56*^-/-^ cells

Having found out that LRRC56 was an IFT cargo, we next sought to explain why it was required for the addition of ODA only to the distal part of the flagellum. The first hypothesis consists of a differential dependence along the length of the flagellum. One strong candidate for this is the docking complex since it is made of a proximal and a distal one in trypanosomes (Edwards *et al*., 2018), but not in *Chlamydomonas* where its composition is homogeneous from base to tip of the axoneme (Owa *et al*., 2014). We first examined whether LRRC56 absence had an impact on the flagellar localization of the distal docking complex members dDC1 and dDC2. Each of the proteins was endogenously tagged with the mNeonGreen (mNG) fluorescent marker in wild-type and *lrrc56*^-/-^ cells. In control cells, live fluorescence imaging revealed that both mNG::dDC1 and mNG::dDC2 distribute to the distal part of the flagella, no matter the phase of construction (Figure 2A), exactly as described (Edwards *et al*., 2018). By contrast, the signal intensity for mNG::dDC1 and mNG::dDC2 was significantly reduced and both proteins often failed to reach the tip of the flagellum in *lrrc56*^-/-^ cells (Figure 2B, white arrows). This pattern is consistent with the absence of outer dynein arms at the distal end of the flagellum reported in *lrrc56*^-/-^ cells (Bonnefoy *et al*., 2018). This result suggests that the cause of the absence of ODA at the distal end is the lack of the distal docking complex.

**Figure 2:**
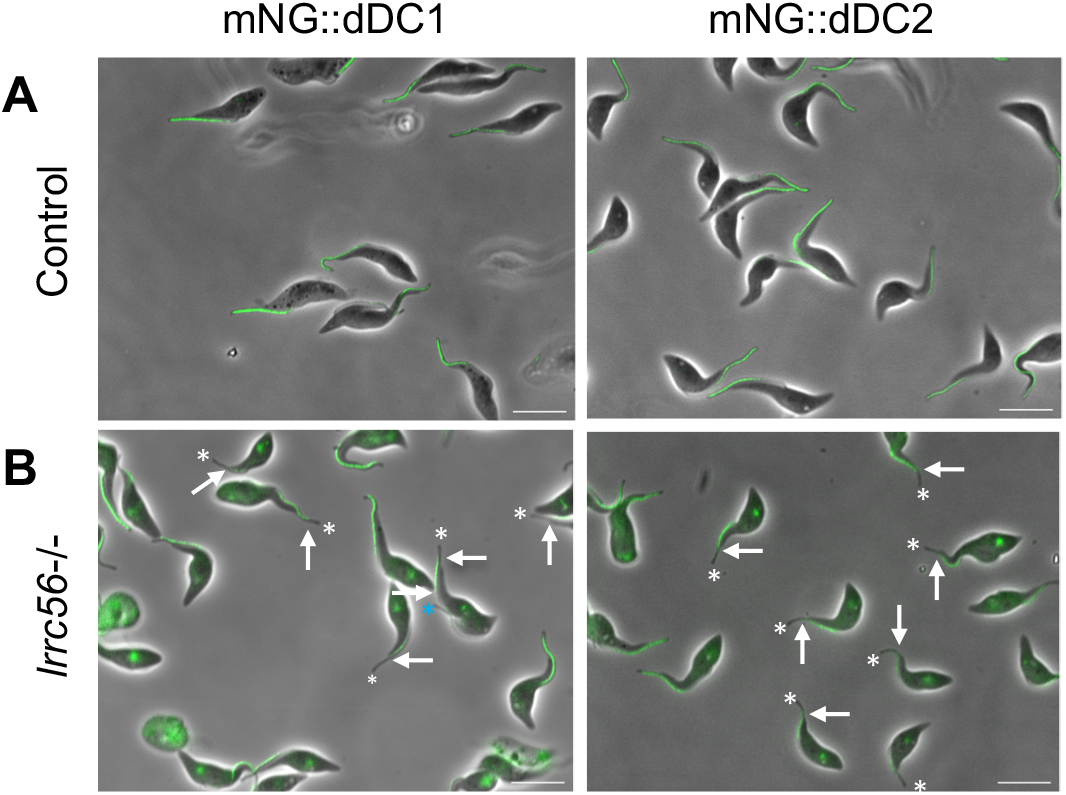
The dDC proteins are missing from the distal part of the flagellum in *lrrc56*^-/-^ cells. *In situ* tagging of dDC1 or dDC2 with mNG was performed in wild-type (A) or *lrrc56*^-/-^ cells (B). In control cells, both mNG::dDC1 and mNG::dDC2 are only found in the distal half portion of the axoneme as expected (A). By contrast, in many *lrrc56*^-/-^ cells, mNG::dDC1 or mNG::dDC2 fluorescence signals are missing from the very distal portion of the axoneme (B), The white arrows show the end of the fluorescent signal. The white and blue asterisks show the end of the mature flagellum (OF) and new flagellum (NF) respectively. Scale bar: 10 *µ*m.

Curiously, this phenotype of the absence of dDC1/2 proteins in *lrrc56*^-/-^ trypanosomes is observed in only about half of cells (white arrows, Figure 3A) while the others harbour complete or nearly complete dDC1/2 staining (red arrows, Figure 3A). To understand the reasons for this heterogeneity, we looked for a possible relationship between flagellum age and maturation (Bertiaux *et al*., 2018b). Trypanosomes that are about to divide possess two flagella, the new one that is under construction and the old one that has been assembled at least one generation before (Sherwin and Gull, 1989). Whilst the mNG::dDC2 signal almost reached the tip of the old flagellum in biflagellated cells (Figure 3B, yellow arrows), the protein is barely detected in the new flagellum, where it fails to reach the distal extremity (white arrows, Figure 3B).

**Figure 3:**
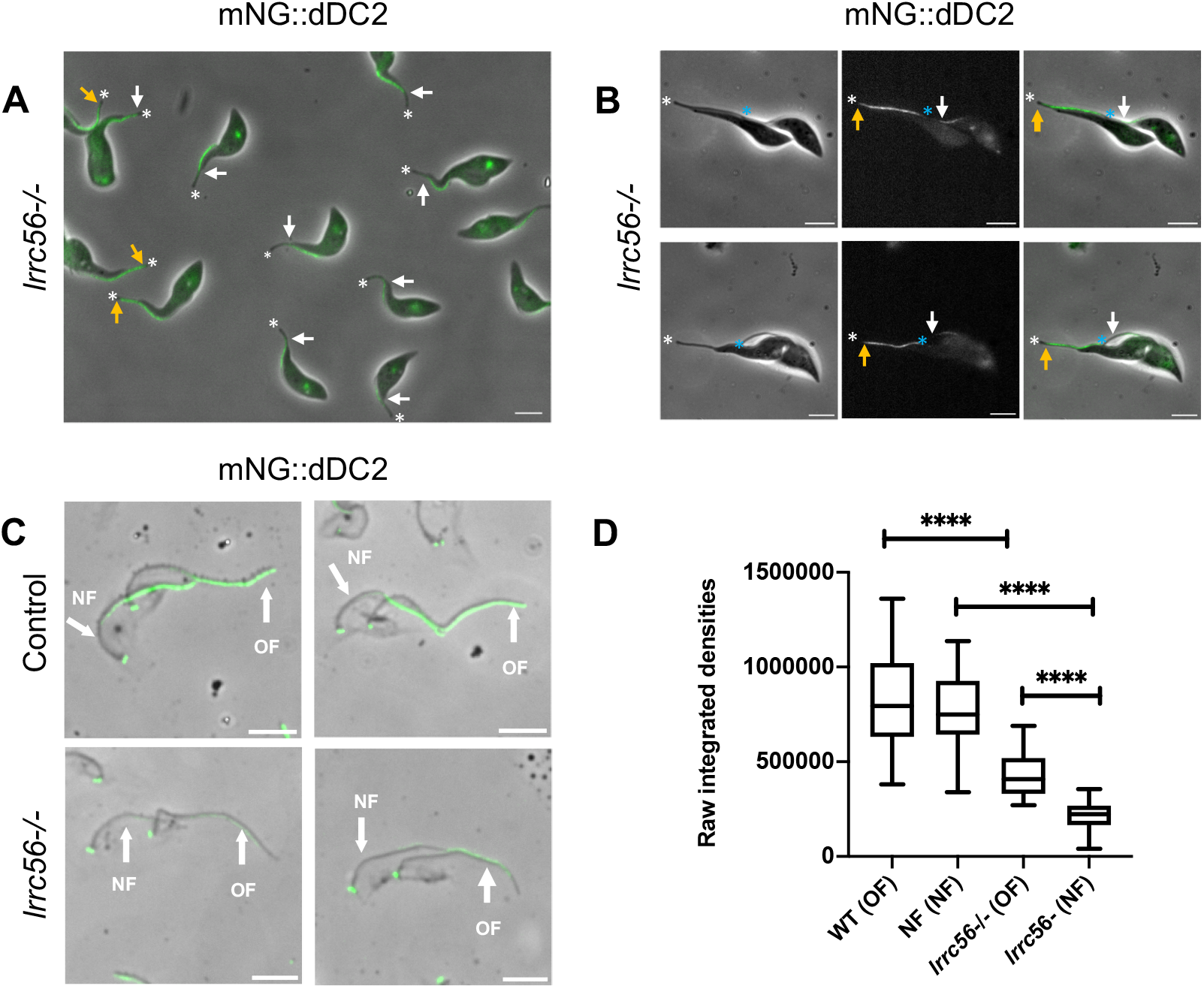
Reduced presence of mNG::dDC2 in *lrrc56*^-/-^ cells is mostly encountered in flagella under construction. A) Distal absence of mNG::dDC2 signal in live *lrrc56*^-/-^ cell line is observed in about half of the cells (indicated with white arrows) while the other cells harbour complete or nearly complete dDC2 flagellar signal except for the tip (yellow arrows). Asterisks show the end of the flagellum. A non-specific signal is also observed at the base of the flagella of tagged mNG::dDC2 cells, exactly as found on control cells (not shown). Scale bars: 5 *µ*m B) In live cells with two flagella about to divide, a nearly complete mNG::dDC2 signal (end of the signal, yellow arrows) is observed in the mature flagellum while in the new flagellum, the signal for mNG::dDC2 (end of the signal, white arrows) is limited to a short, central portion. The white and blue asterisks show the end of the mature flagellum (OF) and new flagellum (NF) respectively. Scale bars: 5 *µ*m (C) In detergent-extracted *lrrc56*^-/-^ cytoskeletons analysed by IFA using an anti-mNG antibody, the absence of mNG::dDC2 is observed at the distal tip of both the old and the new flagellum (white arrows), with reduced signal intensity compared to controls. Images are normalized. Scale bar: 5 *µ*m (D). Integrated density quantification using ImageJ of the total mNG::DC2 signal intensity along flagella in control and *lrrc56*^-/-^ cells revealed a 2-fold decrease in the new flagellum (NF) compared to the old flagellum (OF). 29 cells at 2K2N stage were analysed for both WT and *lrrc56*^-/-^. P-value for each paired data variation was <0.0001 (****).

To evaluate if the mNG::dDC2 protein was correctly attached to the axoneme, cytoskeletons were extracted with detergent, a procedure that removes all soluble material (Robinson *et al*., 1991). In control cells, the mNG::dDC2 protein was tightly associated with the axoneme at its expected distal location in both old and new flagella (Figure 3C, top panels). The low amount of mNG::dDC2 detected by IFA in *lrrc56^-/-^*cells remained associated with the cytoskeleton (Figure 3C, bottom panels), showing that it was properly anchored to microtubules. Since this signal looked weaker compared to controls, its intensity was quantified. No significant differences were seen between old and new flagella of the control cell line. By contrast, a 2-fold reduction of the mNG::dDC2 signal was observed in old flagella of *lrrc56^-/-^* cells compared to controls and a 4-fold reduction in new flagella (Figure 3D).

### Redistribution of the proximal docking complex proteins in *lrrc56*^-/-^ cells

When the distal docking complex is absent upon inducible RNAi knockdown of any of its components, members of the proximal complex (mNG::pDC1 and mNG::pDC2) remain proximal but both components spread toward the distal end, albeit not totally (Edwards *et al*., 2018). We, therefore, wondered whether their localisation was also affected in *lrrc56*^-/-^ cells given the partial depletion of distal docking complex proteins. We have explored this using control or *lrrc56*^-/-^ cells expressing mNG::pDC1 following *in situ* tagging. In control cells, we observed the previously described proximal localization of pDC1 in the first half of the flagellum as expected (Figure 4A). By contrast, the mNG::pDC1 signal extended on much longer portions of the flagellum in *lrrc56*^-/-^ cells, sometimes even reaching the tip of the organelle (Figure 4B).

**Figure 4.**
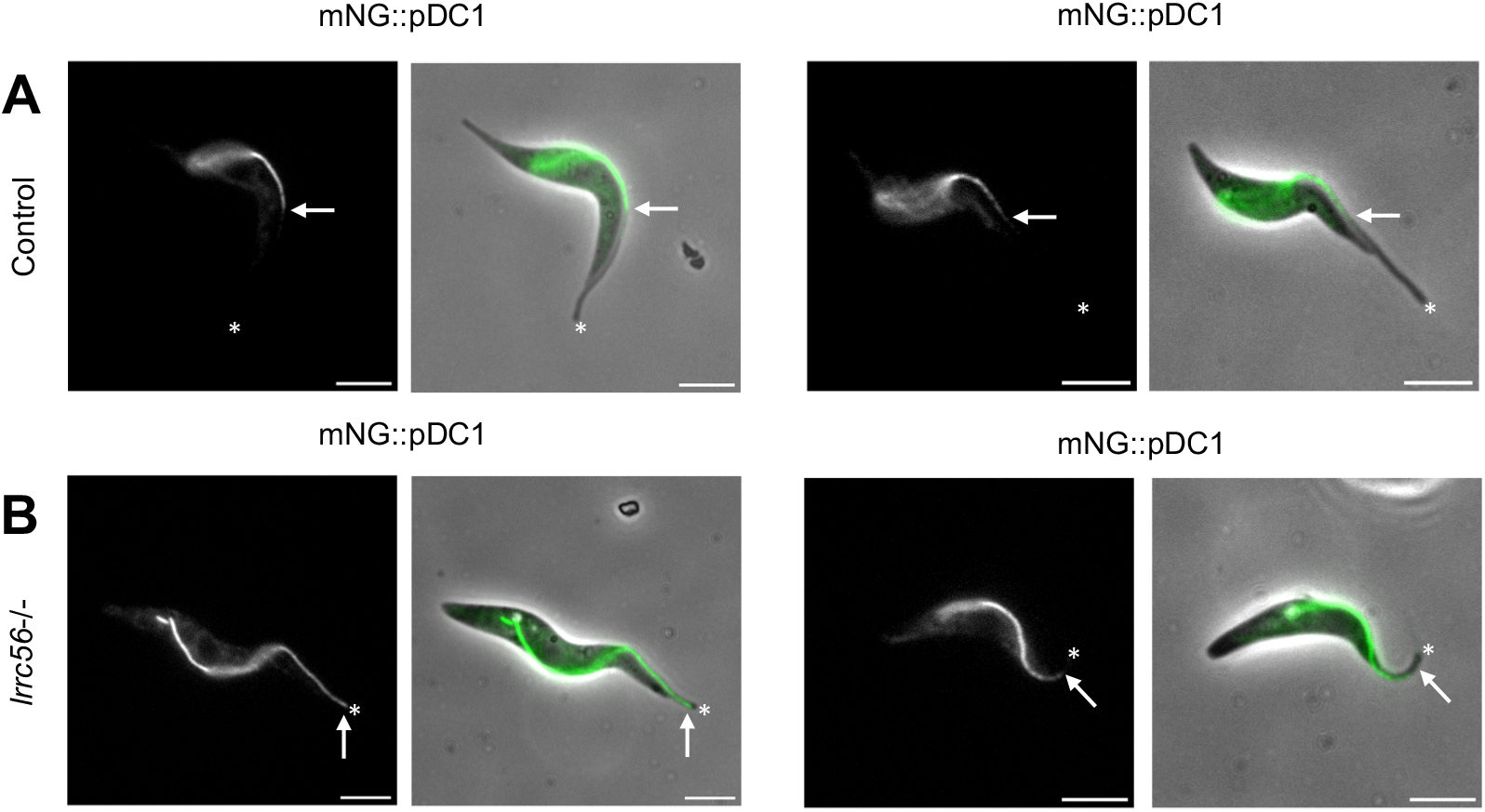
The proximal docking complex covers a much longer portion of the flagellum in *lrrc56*^-/-^ cells. (A, B) *In situ* tagging of pDC1 with mNG in wild-type (A) or *lrrc56^-/-^* cells (B). The mNG::pDC1 signal (green) was merged with the phase contrast image. mNG::pDC1 signal alone is shown at the left of each merged image. The proximal docking complex protein is distributed in the proximal portion of the axoneme (about half-length) in the wild-type background (the white arrows show the end of the fluorescent signal), as expected (A). The distal end of the flagella is shown by a white asterisk. By contrast, the mNG::pDC1 signal extends toward the distal region of the flagellum in *lrrc56*^-/-^ cells (white arrows) (B), sometimes even reaching the distal tip (left panel). Scale bars: 5 *µ*m.

### Dynein arms distribution relies on the proximal DC in *lrrc56*^-/-^ cells

Previous work revealed that the proximal and distal docking complexes occupy roughly each opposing half of the axoneme (Edwards *et al*., 2018). We confirmed this result in cell lines expressing mNG::dDC2 or tdT::pDC1, with measurements showing that each protein covers about 60% of the length of both new and old flagella (Figure 5A). In the middle portion of the flagellum, a decreased signal was observed for each, supporting that distal and proximal docking complex proteins compete for the binding sites in this region of the axoneme (Edwards *et al*., 2018). In *lrrc56*^-/-^ cells, the mNG::dDC2 signal covered only 47% of the length of the old flagellum (Figure 5B) and dropped to barely 26% in new flagella (Figure 5B). By contrast, signals for the proximal docking component expanded to 72% and close to 90% of the total length in new and old flagella, respectively (Figure 5B). To explore whether this redistribution was associated with outer dynein arm docking in *lrrc56*^-/-^ trypanosomes cells, cells were independently stained with the anti-DNAI1 antibody (dynein arm intermediate chain 1, also known as IC78) (Branche *et al*., 2006) that recognizes an essential structural component of the ODA, as previously described (Duquesnoy *et al*., 2009). The distribution of DNAI1 and tdT::pDC1 signals are almost similar in both mature and growing flagella (Figure 5B). This redistribution of the proximal docking complex along the axoneme is quite probably filling progressively the unoccupied distal docking sites, explaining the discrepancies between distal loss of dDCs proteins and more extended axonemal association of ODAs mostly on the old flagellum.

**Figure 5.**
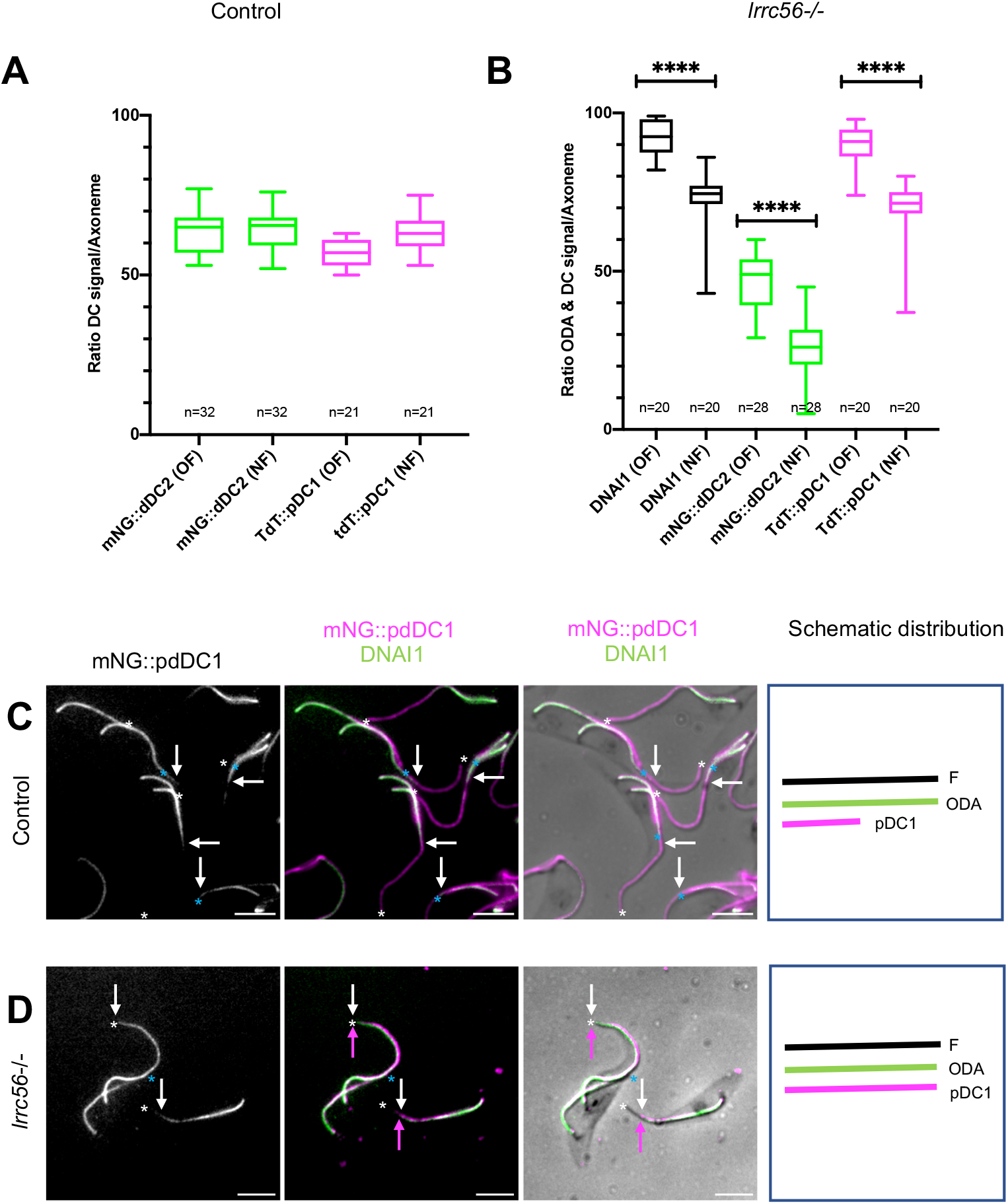
The association of outer dynein arms to the distal portion of the flagellum correlates with the redistribution of the proximal docking complex. (A) distribution of the distal and proximal DCs in old and new flagella of control cells. (B) Distribution of the outer dynein arms (ODA labelled with DNAI1), proximal (tdT::pDC1) and distal (mNG::dDC2) docking complexes in old (OF, left) and new (NF, right) flagella in *lrrc56*^-/-^ cells, showing a correlation between DNAI1 axonemal location and pDC1 redistribution along the *lrrc56*^-/-^ axoneme. The axonemal length was measured using a phase contrast image of the flagellum. P-value for each paired data variation was <0.0001 (****). (C-D) Cytoskeletons of control (C) or *lrrc56*^-/-^ cells (D) expressing mNG::pDC1 and mNG::pDC1 showing localisation of mNG::pDC1 (green) or DNAI1 (magenta). (D) In *lrrc56*^-/-^ cells, the mNG::pDC1 signal extends toward the distal region of *lrrc56*^-/-^ cells (white arrow) correlating to the extent of DNAI1 localisation. The end of DNAI1 localisation is shown by a magenta arrow in the *lrrc56*^-/-^ axoneme. The white and blue asterisks show the end of the monoflagellated and mature flagellum (and new flagellum respectively. Scale bars: 5 *µ*m. Schematic representation of outer dynein arms (ODA) and pDC1 distribution in control and *lrrc56*-/- mature flagellum (F) is shown at the right of panel C-D.

### Dynein arm assembly can be restored in *lrrc56^-/-^* cells

The above results show that the absence of dynein arms at the distal part of growing flagella is related to the molecular heterogeneity of the flagellum along its length, with interdependence between LRRC56 and the proteins of the distal docking complex. However, the fact that ODA absence is less pronounced in mature flagella indicates that a timing issue could also be at stake in the case of different growth rates for axoneme elongation and ODA addition in *lrrc56^-/-^*cells.

To evaluate this possibility, we took advantage of the fact that procyclic trypanosomes assemble their new flagellum in a strictly controlled cell cycle manner, undergoing cytokinesis when the ratio between new and old flagella is only ∼80%, with the rest of the construction taking place after cell division (Sherwin and Gull, 1989; Robinson *et al*., 1991; Farr and Gull, 2012; Bertiaux *et al*., 2018b). We have chemically blocked cell division in *lrrc56^-/-^* cells and control cells expressing mNG::dDC2, mNG::pDC1 or tdT::pDC1 using 10 mM teniposide, an inhibitor of topoisomerase II that interferes with mitochondrial DNA segregation, but not basal body duplication nor flagellum elongation (Robinson and Gull, 1991). This inhibition of cell division allows the completion of the assembly of the new flagellum that reaches the same length as the old one (Bertiaux *et al*., 2018b; Atkins *et al*., 2021). Detergent-extracted cytoskeletons were analysed by IFA using the anti-DNAI1 antibody. Flagellum length was measured and the ratio between old and new flagella was calculated (Figure 6A). In wild-type cells, new flagella have a length at ∼85% of their old counterpart (Figure 6A) and that value climbs close to 100% when cell division is inhibited (Figure 6A), meaning that the length of the new flagellum has reached the length of the old one as previously reported (Bertiaux *et al*., 2018b). Surprisingly, blockage of cell division barely modified the relative length of the new flagellum in *lrrc56^-/-^* cells that moved from 77% to only 81% of the old flagellum length (Figure 6A), a difference that is not statistically significative (p=0.102). By contrast, the presence of dynein arms along old and new flagella was significantly changed in teniposide-treated *lrrc56^-/-^* cells, with dynein arms covering almost the entire length of the axoneme (Figure 6B). This was especially spectacular for new flagella where DNAI1 covers 91% of the length of the axoneme compared to a bit more than 70% in untreated *lrrc56^-/-^* cells as seen previously in Figure 5B (Figure 6B). Similarly, distribution of pDC1 significantly changed in teniposide-treated cells, becoming identical to the distribution of the outer dyneins arms (Figure 6B-D), These results demonstrate that although LRRC56 is missing, if sufficient time is provided, high-affinity association of ODAs to the axoneme can still be achieved all along the mature flagellum and to near completion in the daughter flagellum, presumably because of the compensation by the proximal docking complex (Figure 6B-D).

**Figure 6.**
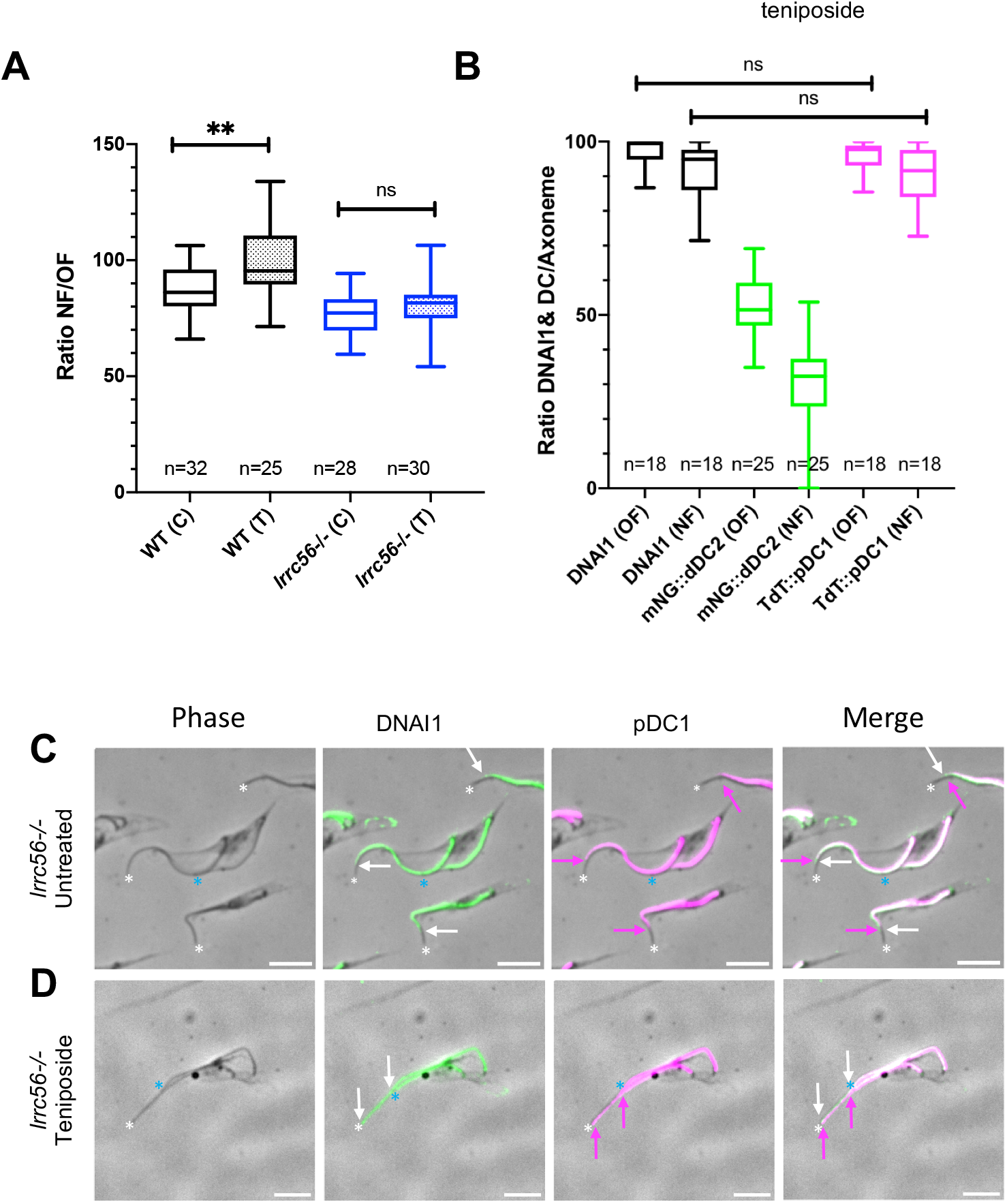
Inhibition of cell division restores most of the outer dynein arms despite the absence of LRRC56. (A) The length of old and new flagella was measured in wild-type and *lrrc56^-/-^*cytoskeletons in the absence or presence of teniposide (T) using phase contrast images of the flagellum. Box plot representing the ratio between the length of the new flagellum (NF) and that of the old flagellum (OF). While the new flagellum reaches the length of the old flagellum in teniposide-treated control cells, inhibition of cell division has little, if any, impact in teniposide-treated *lrrc56^-/-^*cells. (B) Teniposide-treated *lrrc56^-/-^* mNG::dDC2 and *lrrc56^-/-^*mNG::pDC1 cytoskeletons were stained with the anti-DNAI1 antibody as a marker of outer dynein arms and both DNAI1 axonemal distribution and mNG::DC fluorescent signals remaining after fixation were measured. The box plot represents the length proportion of the flagellum that is covered by ODA (DNAI1), proximal (pDC1) and distal (dDC2) docking complex proteins respectively. (C-D) IFA images of cytoskeletons extracted from *lrrc56^-/-^*tdT::pDC1 cells untreated (C) or teniposide treated for 18 hours stained with the anti-DNAI1 antibody (green) and the anti-dsRed (magenta), to detect the tdTomato::pDC1 protein. Distal portions of the flagellum are shown by a white asterisk (monoflagellated or mature flagellum) or blue asterisk (new flagellum). The end of DNAI1 localisation is shown by a magenta arrow in the *lrrc56*^-/-^ axoneme and the end of the tdT::pDC1signal by a white arrow. In *lrrc56*^-/-^ cells treated with teniposide (D), the mNG::pDC1 and DNAI1 signals colocalize and extend toward the distal tip of both mature and new flagellum. Scale bars: 5 *µ*m.

### RNAi knockdown of *dDC2* impairs LRRC56 flagellum localization

The distal docking complex therefore relies on LRRC56 for efficient association with the distal part of the flagellum. To investigate for a reciprocal relationship, we generated inducible RNAi cell lines targeting the *dDC2* gene and expressing mNG::dDC2, mNG::dDC1 or mNG::LRRC56 proteins as reporters. The efficiency of *dDC2* silencing was demonstrated following the endogenous expression of mNG::dDC2. Almost complete loss of the signal (Supplemental Figure S2) was observed upon RNAi induction. In agreement with the prediction that in *C. reinhardtii* the docking complex proteins DC1 & DC2 interact to form a heterodimer (Koutoulis *et al*., 1997; Takada *et al*., 2002), dramatic loss of mNG::dDC1 signal was observed after *dDC2* silencing (Supplemental Figure S2) (Edwards et al., 2018).

Having confirmed the efficiency of dDC2 knockdown, we examined its impact on the presence and localisation of LRRC56 upon *in situ* tagging with mNG. In the absence of RNAi induction, the mNG::LRRC56 protein was found at the distal part of flagella undergoing assembly (Figure 7A), as previously reported for YFP::LRRC56 (Bonnefoy *et al*., 2018). Strikingly, the mNG::LRRC56 flagellar signal was lost in induced *dDC2^RNAi^*cells and the protein was found accumulating in the cytoplasm (Figure 7B) as observed for mNG::dDC1 (Supplemental Figure S2). This result reveals that LRRC56 localisation in the flagellum is dependent upon the distal docking complex proteins. This phenotype is more pronounced compared to the reduction (but not disappearance) of distal docking complex proteins in the absence of LRRC56 (Figures 2 and 3).

**Figure 7.**
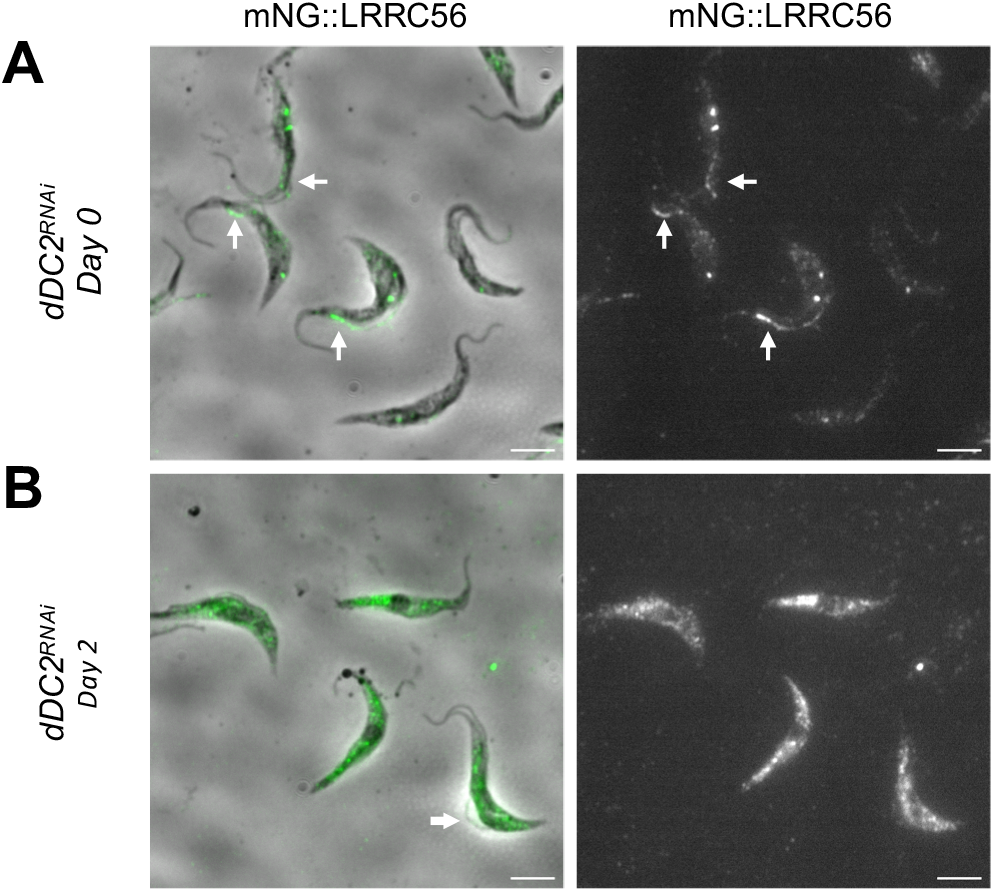
Loss of LRRC56 axonemal localization in the absence of the distal docking complex. Live *dDC2^RNAi^* cells expressing the mNG::LRRC56 protein were observed in control or RNAi-induced conditions. In non-induced cells, the mNG::LRRC56 protein is localized in the distal half portion of growing flagella (A). Following 2 days of dDC2 RNAi induction (B), the mNG::LRRC56 signal is no longer detected in new flagella but the protein is found in cytoplasmic accumulations (B). The images are normalized using ImageJ according to minimum and maximum pixel values. New flagella are indicated by white arrows. Scale bars: 5 *µ*m.

## DISCUSSION

Here, we provide the first direct evidence that LRRC56 traffics in the trypanosome flagellum with similar speed as reported for anterograde IFT. A significant amount of LRRC56 remains stalled with transient episodes of anterograde or retrograde transport, suggesting that LRRC56 would be a transient cargo of IFT trains. Such sequential anterograde progression has also been reported for IFT cargo proteins in *Chlamydomonas* (Wren *et al*., 2013; Craft *et al*., 2015; Dai *et al*., 2018). This observation explains the IFT-dependent localisation previously reported (Bonnefoy *et al*., 2018) and enhances the link between LRRC56 and IFT. However, LRRC56 is able to move within the flagellum independently from IFT. The pattern and relatively fast speed of this movement are unlikely to correspond to diffusion, and the precise mechanisms of these transport events of LRRC56, remain an open question. Phylogenetic distribution of *LRRC56* gene orthologs demonstrated that it is present only in organisms with motile cilia relying on intraflagellar transport for assembly (Desai *et al*., 2015). Expression of the *Drosophila LRRC56* is only observed in chordotonal neurons, the only ones to possess motile cilia assembled by IFT in this organism. Accordingly, it is not detected in the testis where sperm flagellum assembly does not require IFT (Zur Lage *et al*., 2019). Moreover, biochemical analysis has shown that the *Chlamydomonas* LRRC56 protein shares a distribution profile similar to IFT proteins (Desai *et al*., 2015). Finally, pull-down experiments indicate that the human LRRC56 interacts with the IFT88 protein following overexpression in HEK293 cells (Bonnefoy *et al*., 2018).

A second major finding of this study is the unexpected tight interdependence between LRRC56 and the distal docking complex. Depletion of dDC components abolishes LRRC56 presence in the flagellum while knock out of LRRC56 leads to a severe reduction of dDC proteins, hence explaining the absence of ODA only in the distal portion of the flagellum. The apparent more severe impact of the knockdown of distal DC proteins on LRRC56 has to be put in perspective with the significantly different amounts of each protein. LRRC56 was not detected by mass spectrometry of purified flagella, while a large number of peptides derived from dDC components were found: 20 and 14 peptides for dDC1 or 18 and 9 for dDC2 in studies of detergent-extracted flagella or intact flagella, respectively (Broadhead *et al*., 2006; Subota *et al*., 2014). Moreover, the progressive rescue of ODA docking by the proximal docking complex over time solves the problem of the presence of dynein arms on longer portions of flagella as they mature. To explain how LRRC56 could contribute to the proper location of the distal docking complex in the flagellum, we propose that it could function as an IFT adapter protein. Several of these have been reported to facilitate interactions between IFT and candidate cargo molecules (review in Lechtreck, 2022). Such a function could act at the level of entry of dDC components (allowing their passage through the transition zone) or of their transport towards the distal end of the flagellum. Another function could be the unloading of the distal docking complex and perhaps its proper attachment to microtubules.

Edwards *et al*., 2018 have demonstrated that when proteins of the distal complex are absent, those of the proximal docking complex progressively occupy the space left vacant and ensure proper attachment of the dynein arms. Here, the presence of a low amount of dDC proteins means that there is still some competition with pDC proteins which possibly explains why it takes them longer to “colonise” that space. This timing aspect is clearly revealed by the teniposide treatment, which inhibits cell division and hence gives more time for the proximal docking complex to occupy most of the flagellum length.

In *C. reinhardtii*, the transport of ODAs could be visualised directly following the tagging of the IC2/DNAI2 component with the fluorescent mNG protein (Dai *et al*., 2018). Unfortunately, we could not detect the transport either of GFP::DNAI1 or mNG::ODAB (a dynein heavy chain) in growing or mature flagella (Vincensini *et al*., 2018), hence restricting further evaluation of the direct role of LRRC56 in ODA transport. In *Chlamydomonas*, ODAs extracted from wild-type axonemes can stably bind *in vitro* to the axoneme from the *oda8* mutant that locks dynein arms all along the axoneme (Desai *et al*., 2015). By contrast, dynein arms purified from the *oda8* mutant could not attach to wild-type purified axoneme supporting a role for LRRC56 in the maturation of ODAs prior to or during association to the axoneme rather than an involvement in docking complex presence on microtubules (Desai *et al*., 2015).

The unexpected link between LRRC56 and the docking complex that we have highlighted here could be the explanation of the variable impact of LRRC56 mutation or deletion between humans, algae and trypanosomes. Indeed, the composition of the docking complex has evolved differently in these three organisms. The DC was first discovered in *Chlamydomonas* where it is made of at least three proteins called ODA3, ODA1 (two proteins rich in coiled coils) and ODA14 (a calcium-binding protein). While the ODA14 protein seems evolutionary restricted to a few groups (Casey *et al*., 2003), the two other ones are reasonably well conserved in eukaryotes with motile cilia, although they sometimes show extensive divergence (Jerber *et al*., 2014). ODA3 is orthologous to the human CCDC151 and to the trypanosome dDC1 and pDC1 while ODA1 finds orthologous proteins in the human CCDC114 and the trypanosome dDC2 and pDC2. In human cells, two other proteins called ARMC4 and TTC25 are involved in the docking complex and are required for proper outer dynein arm trafficking and docking (Hjeij *et al*., 2014; Wallmeier *et al*., 2016). These have not been found in algae or trypanosomes. Therefore, this significant divergence in the composition of the docking complex could explain the adaptation of its interactions with LRRC56. The trypanosome situation is perhaps the most revealing one, with the distal docking complex requiring LRRC56 for efficient presence in the flagellum while the proximal docking complex seems independent of LRRC56. By contrast, the composition of the docking complex appears homogeneous from base to tip in the flagellum of *Chlamydomonas* (Owa et al. 2014) and in human airway cilia (Hjeij et al., 2014).

No visible impact on dynein arm presence could be detected in the human cilia of a PCD patient with a splicing mutation on the *lrrc56* gene, predicted to encode a non-functional protein (Bonnefoy *et al*., 2018). Recently, another mutation was found in LRRC56, but the cilia of the patient have not been examined (Alasmari *et al*., 2022). If dynein arms are apparently intact, how to explain the altered ciliary beating pattern in this patient? Possibly LRRC56 contributes to more discrete functions for the interactions between dynein arms and the docking complex. For example, LRRC56 might be involved in the proper folding of the dynein arm but not in its attachment. In these conditions, more advanced imaging approaches, such as cryo-electron microscopy could be necessary to evaluate discrete defects, as performed recently to demonstrate the role of tubulin glycylation in the beating of the sperm flagellum (Gadadhar *et al*., 2021).

In conclusion, our results support the hypothesis that the *T. brucei* LRRC56 is a transient IFT cargo protein involved in the binding of the docking complex proteins dDC1 and dDC2 to the distal portion of the axoneme during construction of the new flagellum. The finding of this interaction with the docking complex was unexpected knowing that flagella of the *Chlamydomonas oda8* mutant can bind dynein arms extracted from wild-type cells, suggesting that the docking complex is present and functional in the absence of LRRC56 in this organism. The exhaustive divergence in the composition of the docking complex probably explains the different contributions of LRRC56 to the proper attachment of dynein arms to the docking complex between human, trypanosome and alga axonemes.

## Acknowledgements

E.B. was supported by fellowships from the French National Ministry for Research and Technology (doctoral school CDV515) and from La Fondation pour la Recherche Médicale (FDT20170436836). We thank the UtechS Photonic BioImaging (Imagopole), C2RT, Institut Pasteur, supported by the French National Research Agency (France BioImaging; ANR-10–INSB–04; Investments for the Future), for advice and access to the UltraVIEW VOX system. Research is funded by grants from the ANR (ANR-18-CE13-0014-01), the Fondation pour la Recherche Médicale (FRM-EQU202203014654) and the French Government Investissement d’Avenir programme, Laboratoire d’Excellence “Integrative Biology of Emerging Infectious Diseases” (ANR-10-LABX-62-IBEID).

The authors declare no competing financial interests.

