## Supplemental Figure S2 for "Novel roles for the LRRC56 protein, an IFT cargo protein, in docking of dynein arms in *Trypanosoma brucei*"

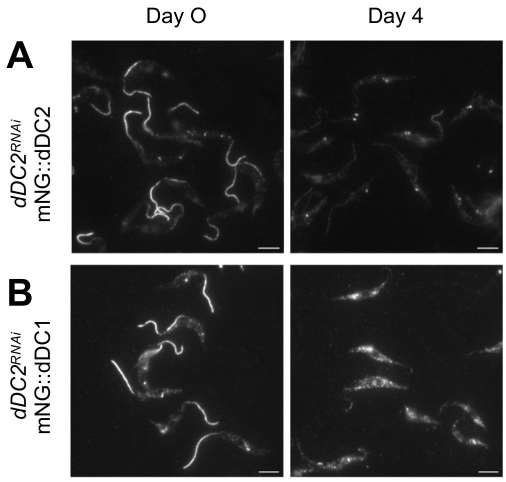


**Supplemental Figure S2. Efficiency of dDC2 knockdown and impact on dDC1.** *dDC2*$^{RNAi}$ cells expressing mNG::dDC2 (A) or mNG::dDC1 proteins (B) were observed by live epifluorescence imaging. In untreated cells (control, left panels), both proteins are localised in the distal half portion of the axoneme, as expected. Following 4 days of *dDC2* RNAi induction (Tet D4, right panels), a nearly complete loss of dDC2 axonemal staining was achieved reflecting knockdown efficiency (A). In *dDC2*$^{RNAi}$-induced cells, mNG::dDC1 was barely associated with the axoneme and most of the fusion protein was found in cytoplasmic accumulations (B). The images are normalized using ImageJ according to minimum and maximum pixel values. Scale bars: 5 $\mu$m.
