## Supplemental Figure S1 for "Novel roles for the LRRC56 protein, an IFT cargo protein, in docking of dynein arms in *Trypanosoma brucei*"

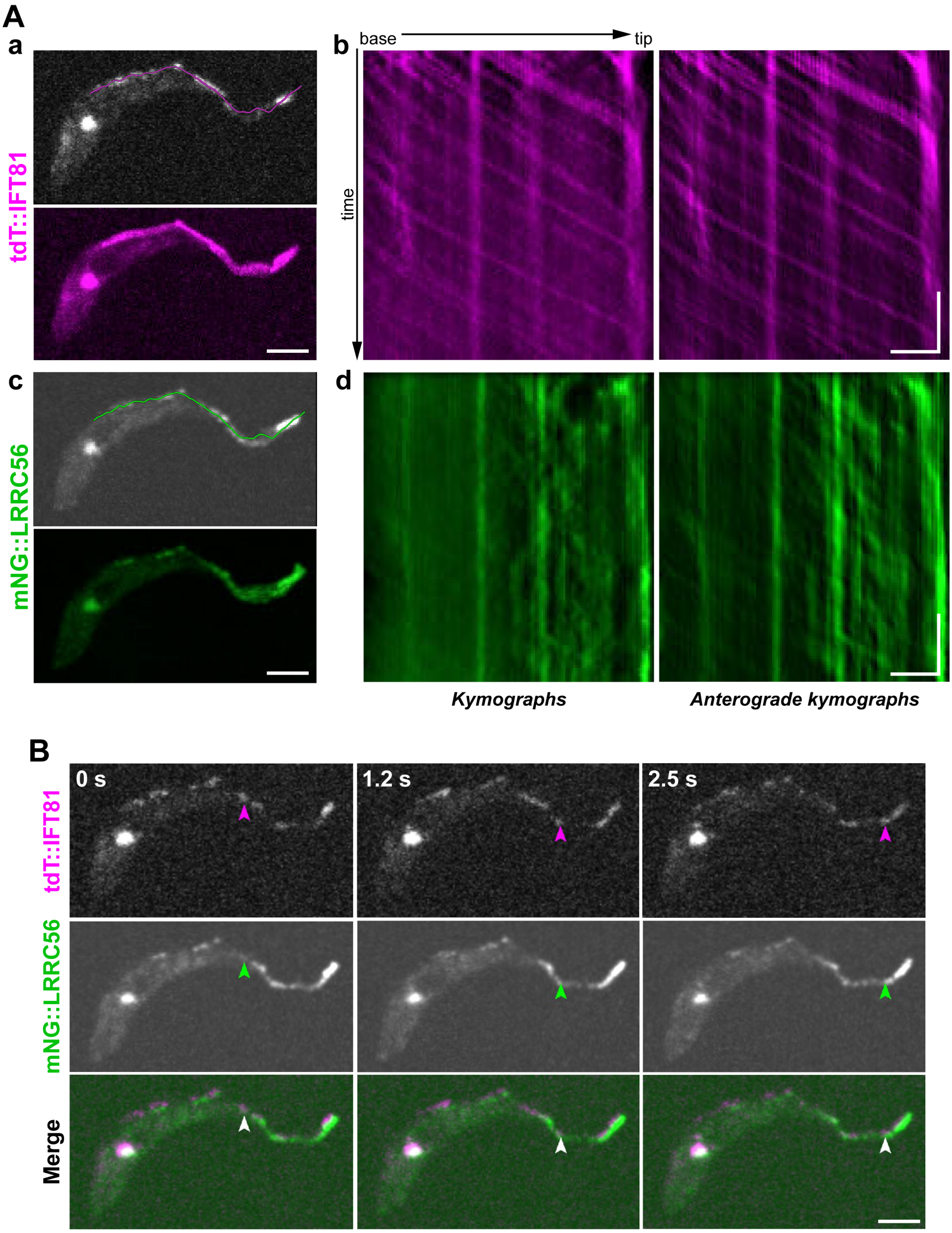


**Figure S1.** (A) Kymograph analysis of a second cell expressing tdT::IFT81 and mNG::LRRC56. (a) A still image of the tdT::IFT81 signal, with the region of interest used to extract the kymographs in magenta (top). 15-second temporal projection of tdT::IFT81 (bottom). Scale bar: 3 $\mu$m. (b) The tdT::IFT81 anterograde kymograph of the cell in (a). Horizontal scale bar: 3 $\mu$m, vertical scale bar: 3 s. (c) A still image of the mNG::LRRC56 signal (top) of the same cell in (a). The region of interest is shown in green. 15-second temporal projection of mNG::LRCC56 (bottom). Scale bar: 3 $\mu$m. (d) The anterograde kymograph of the cell in (c). Horizontal scale bar: 3 $\mu$m, vertical scale bar: 3 s. (B) An anterograde particle containing IFT81 and LRRC56 signals (arrowheads) was observed in still images of the cell in (A) at different time points. The full sequence is in the movie S2. Scale bar: 3 $\mu$m.
